# Holstein Friesian mid-lactation Milk Polar Metabolite Composition in relation to Dietary Fat Composition and Diacylglycerol O-acyltransferase 1 Genotype

**DOI:** 10.1101/303099

**Authors:** Marinus F.W. te Pas, Jacques Vervoort, Leo Kruijt, Mario P.L. Calus, Mari A. Smits

**Author notes:** **Corresponding author**, Marinus F.W. te Pas.

## Abstract

**Background:** The metabolite composition of cow milk is dependent on a large variety of animal associated factors including diet, genotype and gut microbiome composition. The objective of this study was to investigate changes in cow milk polar metabolite composition resulting from dietary and DGAT1 (Diacylglycerol O-acyltransferase 1) genotype perturbations.

**Methods and Results:** Cows were fed a standard diet and a diet supplemented with (poly)unsaturated fatty acids (experimental diet) for ten weeks. Metabolite profiles were determined using ^1^H NMR (1-Hydrogen Nuclear magnetic resonance) technology. The results showed that the diet affected the polar metabolite composition of milk via the metabolism of the cow and via the metabolism of the gut and rumen microbiota. The experimental diet reduced the metabolic rate, especially the energy metabolism and the amino-sugar and amino acid metabolism, of the cows.

**Conclusion:** Our results suggests the DGAT1 genotype affects both the diet related polar metabolite metabolism of the cow as well as that of the rumen microbiota. Milk metabolite levels in animals with more DGAT1 A-alleles were higher than milk metabolite levels in animals with more K-alleles.

## Introduction

The composition of milk is highly variable, even between lactation stages in a single animal [1, 2]. The composition of milk can be influenced by genetic factors of the cow and by the composition of the fodder [3, 4]. The metabolite composition of milk reflects the metabolism of the animal and that of the rumen microbiota [5–7].

The Diacylglycerol O-acyltransferase 1 (DGAT1) gene is known to affect milk yield and its fat composition [8]. The encoded enzyme catalyzes the terminal step of triglyceride synthesis. This reaction is important for triacylglycerol synthesis in the mammary gland, and for the absorption of dietary fats in the intestine [9]. In cattle DGAT1 showed two genotypes differing by one nucleotide substitution resulting in a non-conservative amino acid substitution. The DGAT1 gene has two known genotypes differing for the amino acid 232 and showing two alleles: A (Alanine) and K (Lysine). Mach *et al*. [10] showed that the interaction between the DGAT1 genotype and the cow’s diet regulated the expression level of 30 genes affecting several cellular physiologic and developmental mechanisms and the immune system.

Diet is also known as an important factor in determining milk composition [4–5, 7]. Dietary components regulate both the cow’s metabolism [11–13] and the rumen microbiome metabolism [5–6]. The underlying biological mechanisms probably is regulation of the metabolism via regulation of gene expression [10, 13–15].

Metabolomics is a high throughput methodology to study the metabolite composition of tissues or body fluids. The metabolome consists of both polar and nonpolar molecules of varying molecular weight such as lipids and fatty acids, amino acids and short peptides, but also a mixture of other reaction intermediate products from the metabolism. With ^1^H NMR (1-Hydrogen Nuclear Magnetic Resonance) technology one can detect, identify, and quantify molecules in complex matrices [16–17]. Sundekilde *et al*. [18, 19] previously used ^1^H NMR to evaluate milk metabolite levels in relation to breed and milk properties.

Previously, we reported on the effects of a ten weeks unsaturated fatty acids addition to the diet of mid-lactation dairy cattle [10, 13, 20–22]. These studies showed that adding unsaturated fatty acids to the diet resulted in massive changes in the transcriptome of the mammary gland affecting many cellular metabolic processes and changing the nonpolar milk composition. Furthermore, these studies showed interaction between the diet and the DGAT1 genotype, and the importance of the ruminal metabolism. The impact of these transcriptome changes on the polar metabolite profile remained poorly understood. The objective of this study was to study polar metabolite changes in milk related to unsaturated fatty acids feed composition and DGAT1 genotype during mid lactation in the same animals.

## Materials and Methods

### Animals and experimental conditions

Holstein-Friesian cows in mid lactation were fed diets composed of a standard composition, or with plant oil supplementation (N=7 for each diet; For the DGAT1 genotypes: AA (N=2), AK (N=2), KK (N=3); For all experimental details see Mach *et al*., 2011A, Jacobs *et al*., 2011). In this study we focused on the cows receiving a mixture of rapeseed, soybean and linseed oil together (Table 1, [23]), excluding the animals receiving only one of these oils in an effort to maximize the measurable effects of the dietary differences. All other experimental conditions are as described in [13] because the same samples were used. Milk samples were taken at the end of each feeding period and stored at −25°C until use.

**Table 1.**
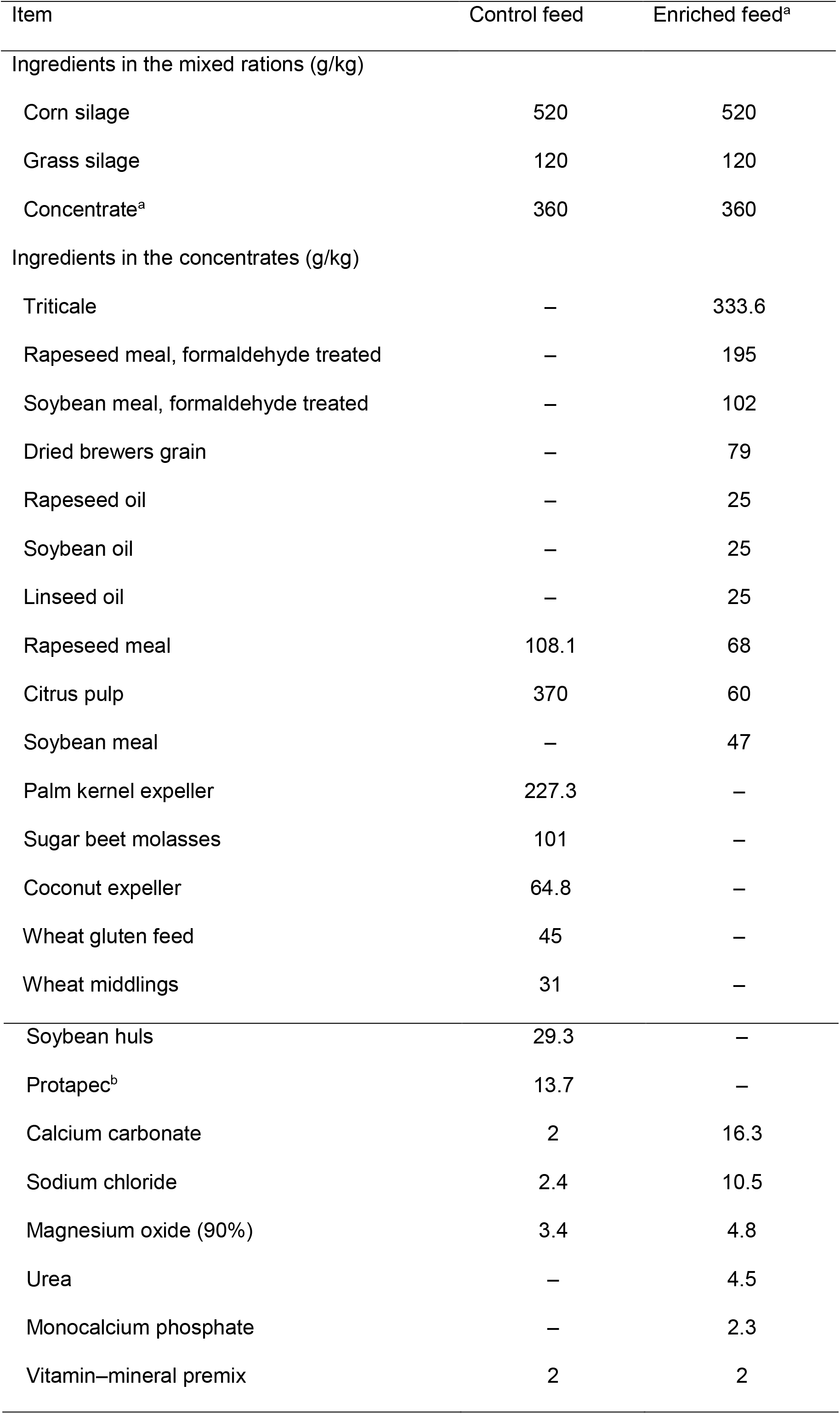
Diet compositions. Control feed is a standard fodder. Enriched feed is standard fodder supplemented with rapeseed, soybean and linseed oil (Data taken from [13, 23]). Mid-lactation cows were fed the control feed and the enriched feed for 10 weeks each

a: Including the concentrate supplied through the automatic feeding station

b: Concentrated potato fruit juice mixed with soybean hulls.

### Diacylglycerol O-acyltransferase 1 (DGAT1) genotyping

Genotyping of DGAT1 gene polymorphism was performed using a TaqMan allelic discrimination method in an Applied Biosystems 7,500 quantitative real-time (qRT) PCR System (Applied Biosystems, Bleiswijk, the Netherlands), as described by [24]. The genotype at the DGAT1 locus was designated homozygous Lys (KK), heterozygous Lys/Ala (KA), or homozygous Ala (AA).

### Nuclear magnetic resonance Metabolomics

Milk samples (300μl) were taken out of the freezer and debris was removed by centrifugation for 10 minutes at 14.000xg at 4 °C. The interphase between pellet and fat layer was collected. To isolate only polar metabolites the non-polar metabolites were removed by chloroform treatment (milk : chloroform: 2:5 vol : vol) The water phases were collected after centrifugation for 10 min, 5000xg at room temperature. Initial ^1^H NMR data showed too broad peaks indicating that additional purification was needed using ultrafiltration with Pall 3K Omega filters (OD003C34) (Pall corporation, Port Washington, NY, USA).

The ^1^H NMR spectra were acquired on a 600Bruker 600MHz Avance III ^1^H NMR spectrometer equipped with a CryoPlatform cryogenic cooling system, a BCU-05 cooling unit, and an ATM automatic tuning and matching unit (Bruker Biospin, Rheinstetten, Germany). The samples were measured in 3-mm ^1^H NMR tubes (Bruker matching system). Measurements were performed at a temperature of 300 K. Baseline corrections and zero alignment were performed manually for all spectra. The spectra were analyzed for background correction using machine-specific manufacturer software. The metabolites were identified using previous knowledge on the ^1^H NMR spectra of cow milk [25–26].

### Statistical analyses and Bioinformatics

The spectra were divided into integrals of 0.04 ppm and the mean signal strength was used as an indicator. Using a T-test approach we studied differences in the signal strength related to (1) feed composition irrespective of DGAT1 genotype generalizing the effect of feed over all animals using an unpaired T-test approach; (2) Because preliminary analysis of the data showed that differences between feeds differed considerably across animals, we studied feed composition per DGAT1 genotype using the paired nature of the experimental design (the same cows received both feeds) using a paired T-test approach, and (3) differences per DGAT1 genotype within one feed using an unpaired T-test approach. A higher stringency ANOVA analysis was performed using the function AOV of the R-package STATS. Two models were used: (1) Test for feed composition effects: y = μ + feed composition + e (where y = phenotype; μ = overall mean; e = residual) (Single-factor ANOVA); (2) Test for feed composition and DGAT1 genotype: y = μ + feed composition + DGAT1 genotype + e (Two-factor, no interaction ANOVA). For the DGAT1 genotype analysis the animals were grouped together per genotype. For all statistics: P<0.05 was considered significant.

Biological functional analysis was done for all significant identified metabolites using available databases on the internet, e.g. KEGG (Kyoto Encyclopedia of Genes and Genomes; http://www.genome.ad.jp/kegg/ [27]) and Gene Ontology (http://geneontology.org/, 28).

## Results

### Identified polar metabolites in the milk of mid-lactation cows

The ^1^H NMR spectra showed 83 identifiable peaks (Table 2), of which the identity of six peaks remained unknown. Since several metabolites were characterized / represented by more than one peak, in total 57 different metabolites were identified. Several of these metabolites are related to each other – e.g. phosphorylated and non-phosphorylated forms. After correction for multiple forms of a metabolite, 49 different polar metabolites were detected in the milk samples. Two of them were identified only as “a sugar”, without further specification. The integrals harboring the unidentified peaks were included in the analyses to evaluate the effects of diet and DGAT1 genotype. Interestingly, the databases indicated that several of the metabolites were derived from bacterial metabolism, e.g. Xantine, N-acetyl-glucosamine, and butyrate.

**Table 2.**
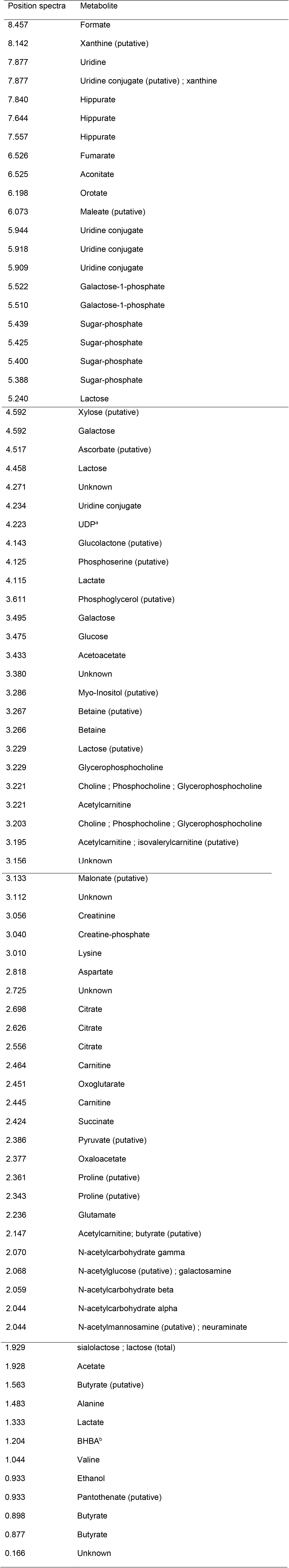
1-Hydrogen Nuclear magnetic resonance detected polar metabolites in Holstein Friesian milk identified by their peak position in the 1-Hydrogen Nuclear magnetic resonance spectra.

a: Uridine diphosphate;

b: Beta-hydroxybutyrate.

### Associations between polar metabolite concentrations in the milk and diet

Table 3 shows the associations between identified metabolites and diet. The Supplementary Table S1 which is related to Table 3, shows all associations including those of the unidentified integrals of the ^1^H NMR spectra. More specifically, Table 3 shows the association between the measured milk polar metabolite levels and the feed composition of the two diets with different fat composition. If more than one *P*-value is expressed in Table 3, the associations stretches over more than one integral and the integrals were taken together. After the first analysis, we included a high-stringent analysis to confirm the associations. As expected, the number of associations in this high-stringency analysis was lower than in the first analysis, but integrals that related to the same metabolites were found in both analyses. Remarkably, many of the observed associations differed between the animals indicating an animal-specific component in the associations.

**Table 3.**
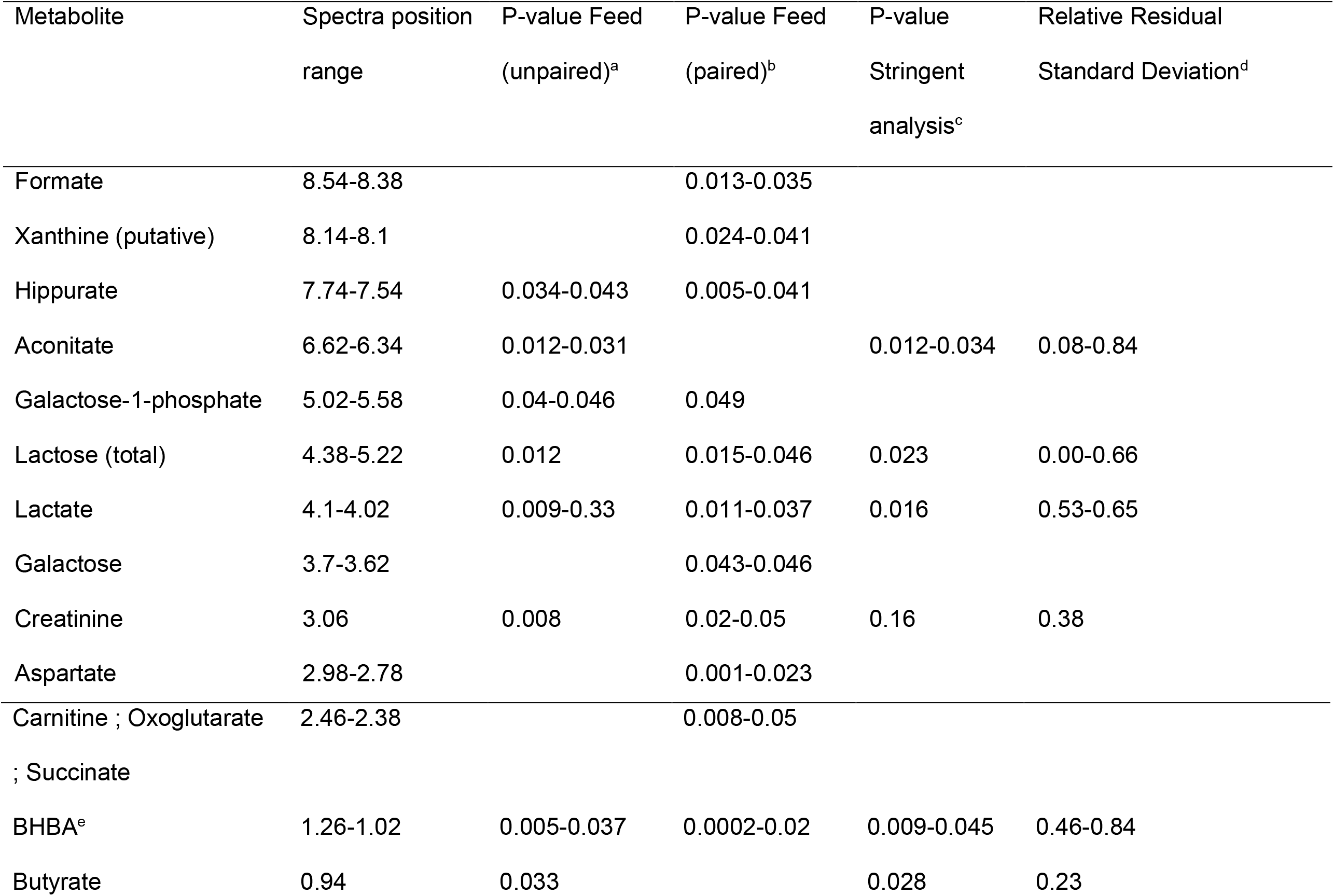
Association between mid-lactation cow milk polar metabolite levels detected and quantified by 1-Hydrogen Nuclear magnetic resonance spectroscopy and diet with different fat composition. Mid-lactation cows were fed a standard fodder or a standard fodder supplemented with rapeseed, soybean, and linseed oil for 10 weeks each

a: Analyzed the dietary groups;

b: Analyzed the metabolite levels between the two diets within each animal individually;

c: High stringent statistical analysis using model 1;

d: The control diet was set to 1.00;

e: Beta-hydroxybutyrate

### Effects of Diacylglycerol O-acyltransferase 1 genotype

Using the same ^1^H NMR measurements the effects of the DGAT1 gene genotypes on the polar metabolite levels were studied. Table 4 and Supplementary Table S2 which is related to Table 4 show the associations between the two DGAT1 genotypes and the content levels of polar metabolites. More specifically, Table 4 shows the association between the measured milk polar metabolite levels differently expression among the three DGAT1 genotypes. Differences between the genotypes for polar metabolite levels in the milk were observed for both diets, but more for the experimental diet than the standard diet. The association between polar metabolites and DGAT1 genotype did not reveal any animal effects. Please note that the high stringent analysis indicated associations for a number of the associations found also with the low stringent analysis, but this high stringent analysis was unable to specify the genotypes.

**Table 4.**
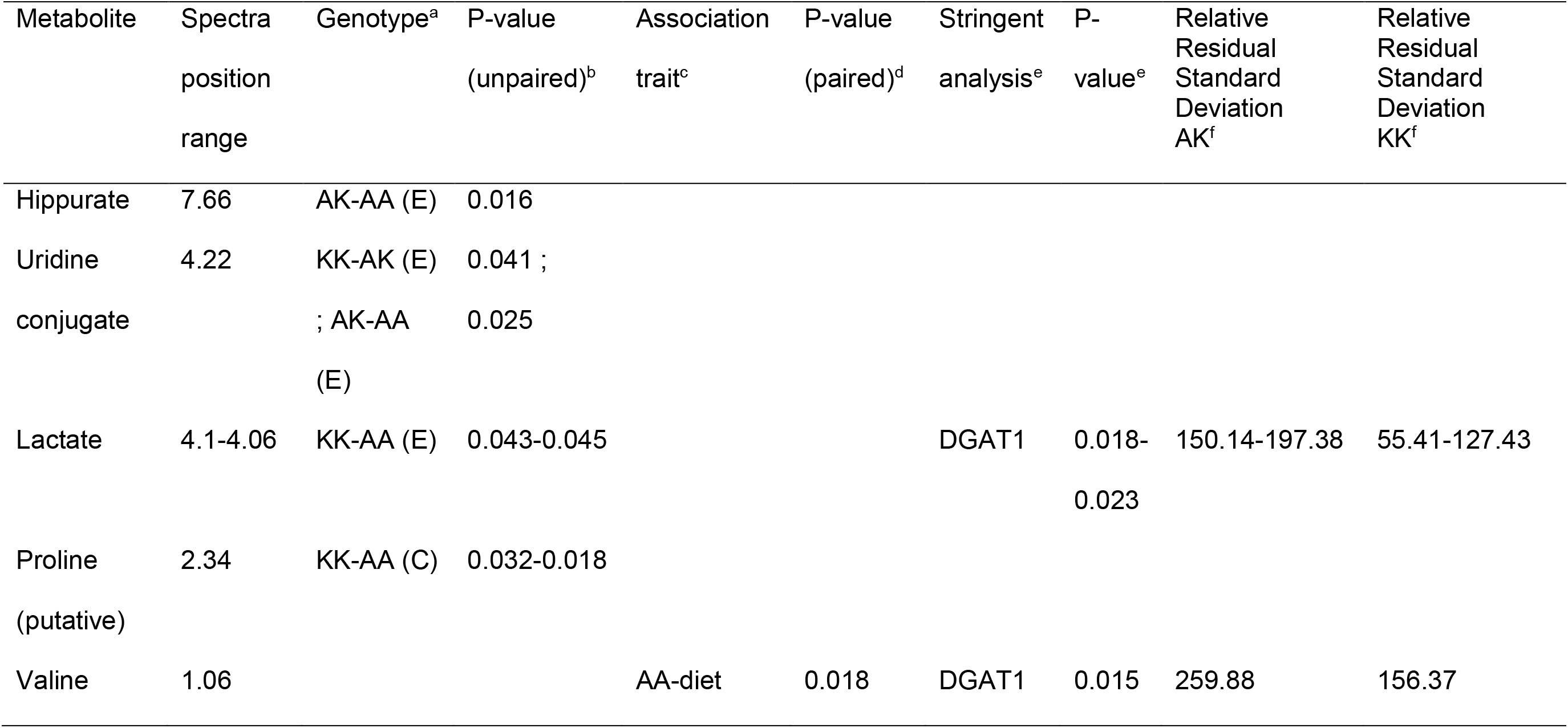
Association between mid-lactation cow milk polar metabolite levels detected and quantified by 1-Hydrogen Nuclear magnetic resonance spectroscopy and diet with different fat composition. Mid-lactation cows were fed a standard fodder or a standard fodder supplemented with rapeseed, soybean, and linseed oil for 10 weeks each. The analysis compared the milk metabolite levels between Diacylglycerol O-acyltransferase 1 genotypes and diets

a: DGAT1 genotypes with significant different milk polar metabolite levels, E: experimental diet (i.e. supplemented with oils), C: Control diet (i.e. standard fodder);

b: Analyzed the dietary groups;

c: Association between DGAT1 genotype and diet;

d: Analyzed the metabolite levels between the two diets within each animal individually;

e: High stringent statistical analysis using model 2;

f: The value of the AA genotype was set to 1.00.

### Effect of Diacylglycerol O-acyltransferase 1 genotype on dietary-induced milk polar metabolite levels

In a next step, we studied the effect of the DGAT1 genotype on dietary-induced milk polar metabolite concentrations in order to investigate the interaction between the diet and the DGAT1 genotype effects on the content levels of polar metabolites. Table 5 and Supplementary Table S3 which is related to Table 5 show that the DGAT1 A-allele displays more interactions with the diet than the K-allele. More specifically, Table 5 shows the association between the measured milk polar metabolite levels differently expression among the three DGAT1 genotypes specifically related to the interaction between the feed composition and the DGAT1 genotypes. Furthermore, the interaction between the diet and the DGAT1 genotype differed considerably between animals. The number of high-stringent analysis associations was low in this analysis, which suggests that the importance of the interaction between the diet and the DGAT1 genotype is relatively low.

**Table 5.**
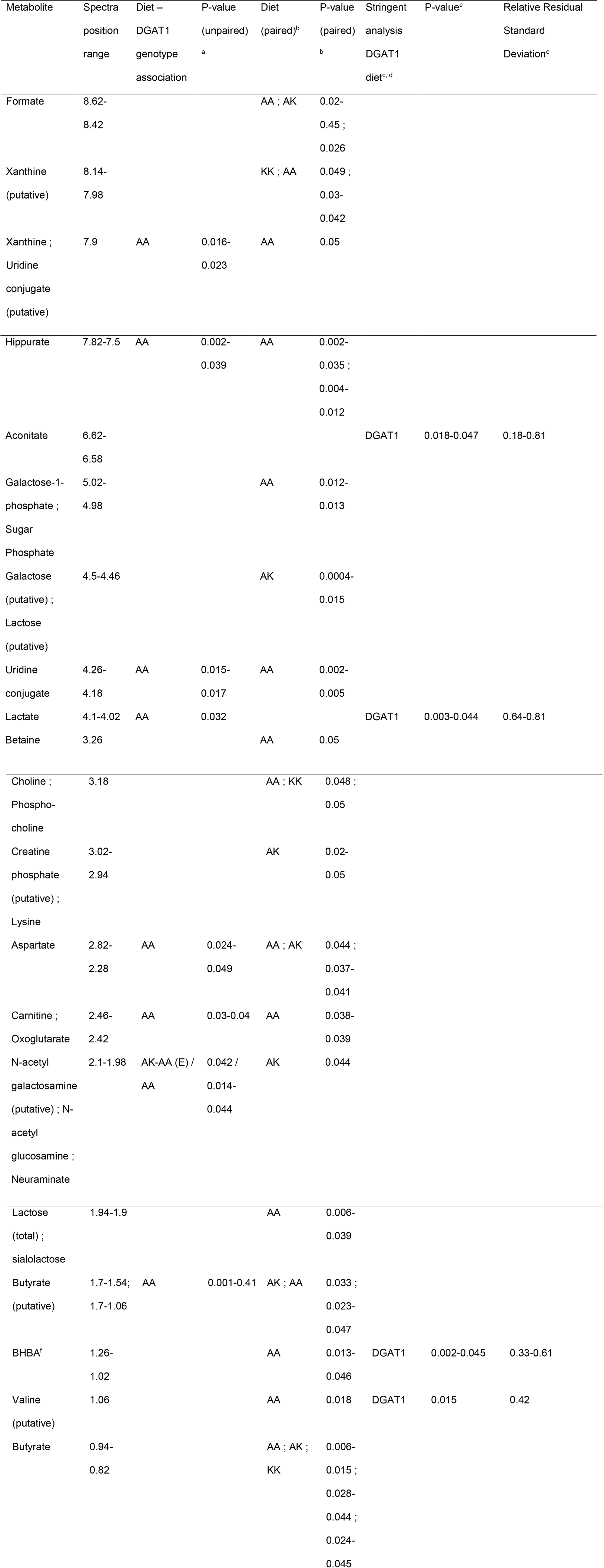
Association between mid-lactation cow milk polar metabolite levels detected and quantified by 1-Hydrogen Nuclear magnetic resonance spectroscopy and diet with different fat composition. Mid-lactation cows were fed a standard fodder or a standard fodder supplemented with rapeseed, soybean, and linseed oil for 10 weeks each. The analysis compared the interaction between milk metabolite levels and diet-Diacylglycerol O-acyltransferase 1 genotype.

a: Analyzed the dietary groups;

b: Analyzed the metabolite levels between the two diets within each animal individually;

c: High stringent statistical analysis using model 2;

d: DGAT1: Diacylglycerol O-acyltransferase 1 e: The control diet was set to 1.00;

f: Betahydroxybutyrate

## Discussion

### Dietary fats and milk polar metabolites

Previously it was demonstrated that dietary fatty acids affect the composition of non polar metabolites in cow milk [5–7]. In this study we have identified a number of polar metabolites in cow milk that are related to dietary fat consumption. Together, this indicates that dietary fatty acids influence the entire cow’s metabolism in such a way that it also affects a wide range of milk traits. This is in agreement with suggestions from several earlier studies [12], and [13, 20, 22] using these same samples.

### Dietary and genetic influence on milk polar metabolite concentration

Feeding extra unsaturated fatty acids (UFA) to the cows was associated with changes in the concentration of several polar metabolites in the milk which were not directly related to the UFA. Thus, it may be concluded that the changed milk composition is related to changed metabolism – either of the cow, or of the gut or rumen bacteria of the cow. Indeed, the results show examples of both. It has been reported that fatty acids may induce or inhibit the expression of many different gene (sets) and thereby change the metabolism of animals [10–11, 13–14].

### General metabolism

In general, the concentration of most polar metabolites was lower in the milk of the animals with the UFA-supplemented diet as compared to the standard diet. Thus, UFA supplementation reduced the metabolism for polar metabolites of the animal. The UFA molecules are involved in many diverse metabolic pathways including energy metabolism. It is known that high energy intake affects animal metabolism rates [7]. However, importantly, lactose (milk sugar) showed increased levels in the milk of UFA supplemented cows. This suggests that the metabolic effects are specific and not general.

### Energy metabolism and Nitrogen metabolism / urea cycle

The levels of ten polar metabolites with association to the dietary composition related to energy metabolism: succinate, formate, aconitate, lactate, galactose-1-phosphate, creatinine, carnitine, oxoglutarate, BHBA (beta-hydroxybutyrate), and butyrate. The concentrations of these metabolites were lower in the diet supplemented with the UFA suggesting that the effect of adding UFA to the diet is to reduce the rate of the cow’s energy metabolism. This has been observed before [11, 15, 29]. It should be noted that butyrate is produced by the gut microbiota serving as an energy source for the colon cells metabolism rather than being produced by the cow’s own metabolism. Effects of UFA in feed on ruminal metabolism has been reported before [5].

Hippurate and formate levels in milk were variable between dietary groups and animals, but no clear direction could be found. Aspartate showed reduced levels in the milk of UFA supplemented cows. We conclude that UFA supplementation did affect nitrogen metabolism in the cows, but probably the cow’s genotype may specifically relate to the specific direction of the effect.

### Animal-specific effects – the Diacylglycerol O-acyltransferase 1 genotype of cows

It is intriguing to note that the animals differed considerably for the majority of the associations. This indicate that the reaction of the animals to the diet was very different, so animal specific components such as genotype may be determining factors for dietary responses. The associations between metabolite concentrations and DGAT1 genotype did not show this animal-specific effect. Therefore, we suggest that the genetic component differentiating between animals may be the DGAT1 genotype, a new feature of this genotype. Although the number of cows per DGAT1 genotype is low, different types of analyses consistently indicate the same metabolites with different profiles per DGAT1 genotype. Furthermore, these differences are also consistent with previously reported general DGAT1 genotype effects. Therefore, these results can be trusted. DGAT1 genotype was already known to be related to milk yield and milk fat percentage and composition [8]. These DGAT1 genotype effects may also relate to the cow’s metabolism or gut and rumen microbiome metabolism rate, and may therefore relate to this new feature of the DGAT1 genotype that we describe here. This feature may perfectly fit into the biological function of DGAT1, which regulates the final step before uptake of fat in the intestine [9]. Thus, the two genotypes of the DGAT1 gene may differ in the amount of uptake of fatty acids in the intestine, thereby differentially regulate the animal’s metabolism including energy metabolism leading to different milk yield and composition. Furthermore, Sorensen *et al*. [30] showed DGAT1 genotype effects on DGAT1 activity. This is a further argument that the DGAT1 genotype may also affect milk composition via regulation of the genome activity via regulation of the intestinal fatty acid uptake.

With the exception of Hippurate, valine, and Uridine-conjugate the levels of the metabolites in the milk were higher in animals with more DGAT1 A-alleles than in animals with more K-alleles (i.e. comparing the metabolite levels in the milk of animals with the AA-genotype with the milk metabolite levels of animals with the AK and KK genotypes, and comparing the metabolite levels in the milk of animals with the AK-genotype with the milk metabolite levels of animals with the KK genotype). This suggests that the DGAT1 genotype associates with the rate of metabolic processes. Metabolites showing association between metabolite levels and DGAT1 genotype were involved in energy metabolism and nitrogen metabolism including amino acid and pyrimidine metabolism pathways. On the other hand, proline shows the opposite content pattern in relation to the DGAT1 genotype, so we cannot conclude that amino acid metabolism is higher regulated to the K-allele than the A-allele. High stringency analysis confirmed several of the associations. Allele-specific effects on amino acid metabolism were not previously observed. General genotype-specific effects on amino acid plasma levels, which could be related to amino acid metabolism, have been reported before [31]. Such plasma level variations could affect amino acid composition in the milk, differentiating animals, which we detected in our metabolomics analyses.

Interactions between the effects of diet and the effects of DGAT1 genotype were prominent. Above we noted that most metabolites associated with diet showed lower levels of the metabolites in the milk of animals fed the UFA-supplemented diet as compared to the control diet. However, here it is interesting to note that the opposite content pattern was mainly observed in animals with the AK-DGAT1-genotype. Grisart *et al*. [8] reported the lack of dominance effects for the DGAT1 alleles. Our results indicating the opposite content pattern in the heterozygous animals may relate to this, since both homozygous genotype animals showed the opposite content pattern. Alternatively, this result could indicate again a direct influence of the DGAT1 genotype on cow or rumen bacteria metabolism.

Indeed, the KEGG and other databases used identified several metabolites as compounds related to (unspecified) bacterial metabolism, e.g. Xantine (microbial metabolism in diverse environments) and N-acetyl-glucosamine (bacterial cell wall component), and butyrate (known for gut microbiome metabolism – it is an energy source for the colon epithelial cells). Since these metabolites are exclusively synthesized by bacterial metabolism and cannot be synthesized by a mammalian metabolism, finding these metabolites in the cow’s milk may indicate a direct connection between the cows DGAT1 genotype and the rumen microbiota metabolic rate. An interaction between animal’s genotype and the rumen bacterial population has been observed before [32]. Dietary composition effects on the gut bacterial composition has also been shown before [32–34]. So, this result may relate to both of these effects and may further show the importance of the interaction between the cow’s genotype and the rumen bacterial population metabolic activity. In conclusion, we showed that both diet and DGAT1 genotype affect polar metabolite concentration in cow milk. Specifically, we showed effects on energy metabolism and nitrogen metabolism in the cow, and bacterial metabolism in the gut bacteria. Our data suggest that both cow metabolism and rumen microbiota metabolism are important for the milk polar metabolite concentrations, and that a direct interaction between the cow’s DGAT1-genotype and the activity of the rumen bacterial metabolism is important for the cow’s milk polar metabolite composition. The interaction between diet and DGAT1-genotype also affect the cow’s milk polar metabolite composition.

## ACKNOWLEDGEMENT

The authors thank Dr. R.C.H. de Vos (Plant Research International, Bioscience, Wageningen UR) and Dr. A. Lommen (RIKILT, Wageningen UR) for helpful discussions during the project. The work was funded by a grant from the Dutch ministry EL&I, grant no. KB-15-001-020.01. The funders had no role in study design, data collection and analysis, decision to publish, or preparation of the manuscript.

---

**Supplementary Table 1.**
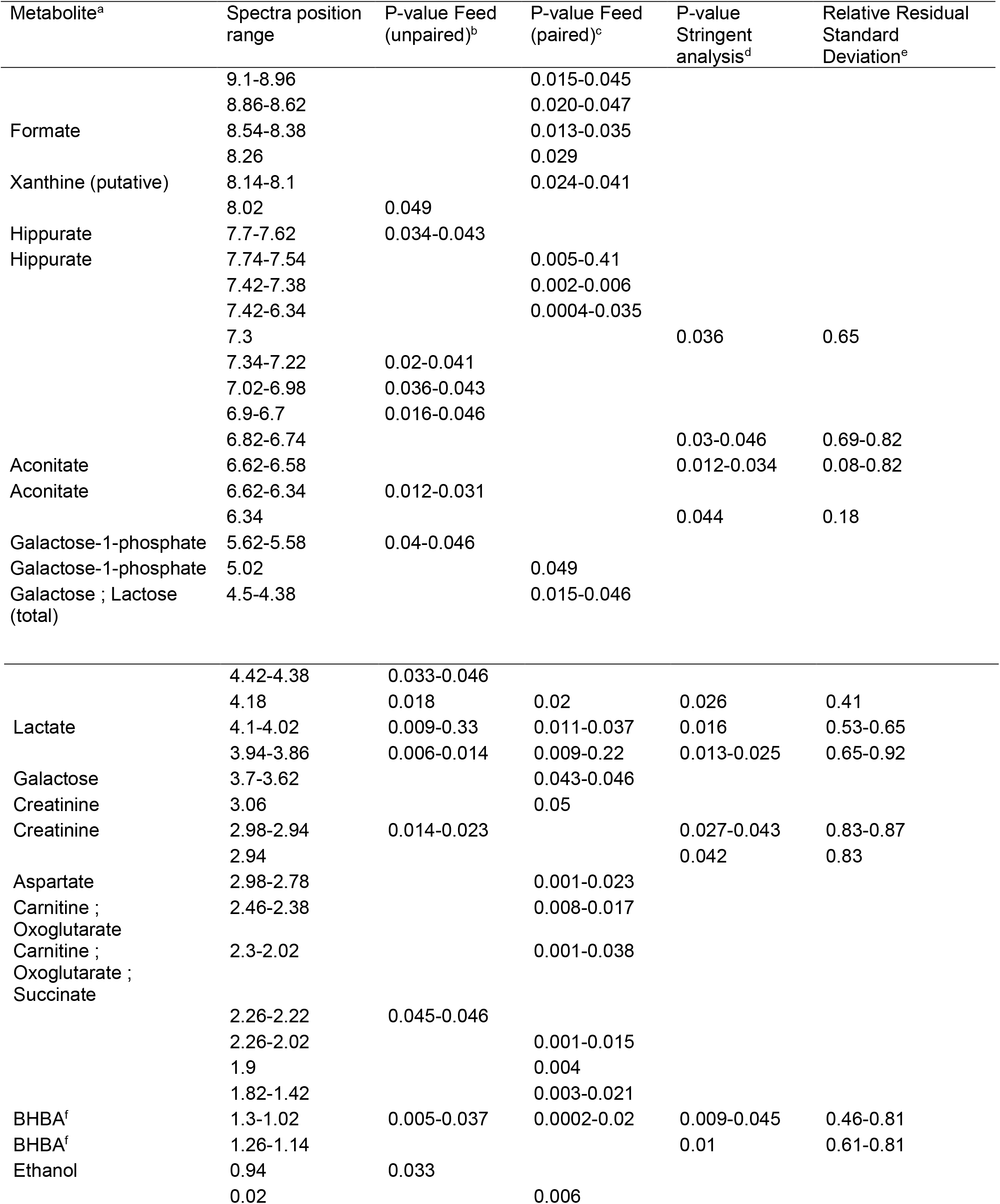
Supplementary Table related to Table 3. Association between mid-lactation cow milk polar metabolite levels detected and quantified by 1-Hydrogen Nuclear magnetic resonance spectroscopy and diet with different fat composition. Mid-lactation cows were fed a standard fodder or a standard fodder supplemented with rapeseed, soybean, and linseed oil for 10 weeks each. This supplementary Table also shows the associations with unidentified integrals of the 1-Hydrogen Nuclear magnetic resonance spectra.

a: Open cells indicate unknown metabolites;

b: Analyzed the dietary groups;

c: Analyzed the metabolite levels between the two diets within each animal individually;

d: High stringent statistical analysis;

e: The control diet was set to 1.00;

f: Beta-hydroxybutyrate

**Supplementary Table 2.**
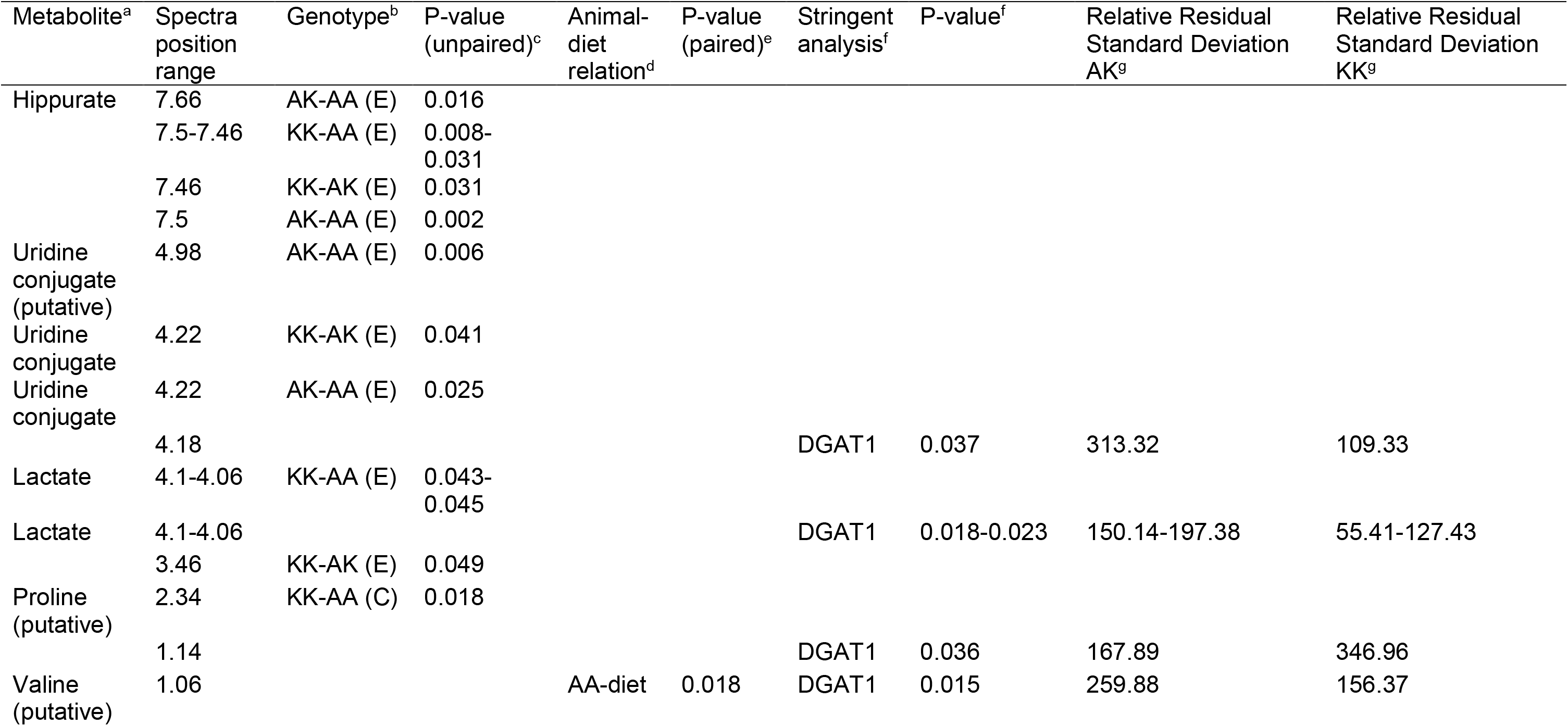
Supplementary Table related to Table 4. Association between mid-lactation cow milk polar metabolite levels detected and quantified by 1-Hydrogen Nuclear magnetic resonance spectroscopy and diet with different fat composition. Mid-lactation cows were fed a standard fodder or a standard fodder supplemented with rapeseed, soybean, and linseed oil for 10 weeks each. The analysis compared the milk metabolite levels between Diacylglycerol O-acyltransferase genotypes and diets. This supplementary Table also shows the associations with unidentified integrals of the 1-Hydrogen Nuclear magnetic resonance spectra.

a: Open cells indicate unknown metabolites;

b: DGAT1 genotypes with significant different milk polar metabolite levels, E: experimental diet (i.e. supplemented with oils), C: Control diet (i.e. standard fodder);

c: Analyzed the dietary groups;

d: Association between DGAT1 genotype and diet;

e: Analyzed the metabolite levels between the two diets within each animal individually;

f: High stringent statistical analysis;

g: The value of the AA genotype was set to 1.00.

**Supplementary Table 3.**
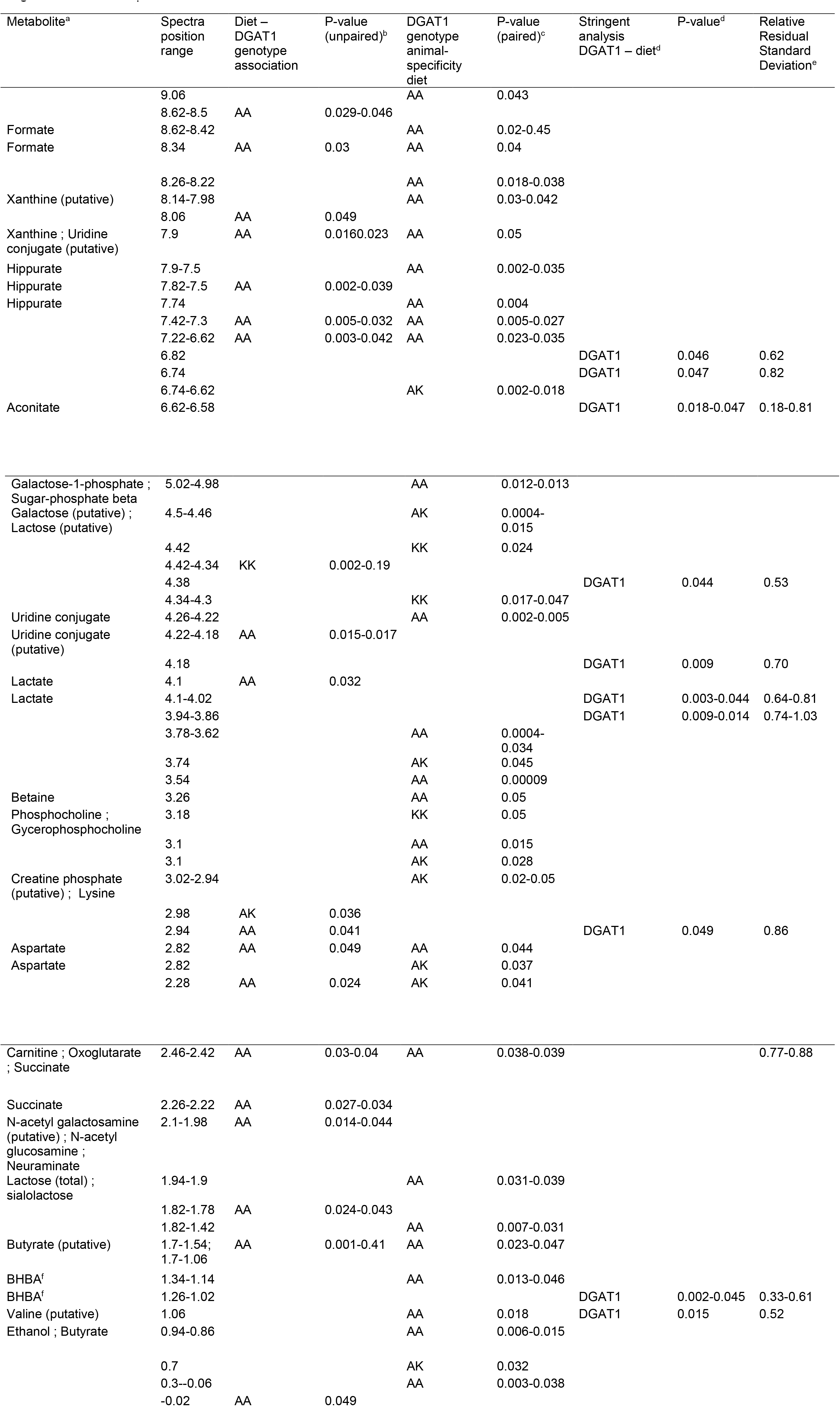
Supplementary Table related to Table 5. Association between mid-lactation cow milk polar metabolite levels detected and quantified by 1-Hydrogen Nuclear magnetic resonance spectroscopy and diet with different fat composition. Mid-lactation cows were fed a standard fodder or a standard fodder supplemented with rapeseed, soybean, and linseed oil for 10 weeks each. The analysis compared the relation between milk metabolite levels and diet-Diacylglycerol O-acyltransferase genotype. This supplementary Table also shows the associations with unidentified integrals of the 1-Hydrogen Nuclear magnetic resonance spectra

a: Open cells indicate unknown metabolites;

b: Analyzed the dietary groups;

c: Analyzed the metabolite levels between the two diets within each animal individually;

d: High stringent statistical analysis;

e: The control diet was set to 1.00;

f: Betahydroxybutyrate

## REFERENCES

1. Kelsey JA, Corl BA, Collier RJ, Bauman DE (2003) The Effect of Breed, Parity, and Stage of Lactation on Conjugated Linoleic Acid (CLA) in Milk Fat from Dairy Cows. J Dairy Sci 86:2588–2597.

2. Stoop WM, Bovenhuis H, Heck JML, van Arendonk JAM (2009) Effect of lactation stage and energy status on milk fat composition of Holstein-Friesian cows. J Dairy Sci 92:1469–1478.

3. Grisart B, Farnir F, Karim L, Cambisano N, Kim J-J, Kvasz A, Mni M, Simon P, Frère J-M, Coppieters W, Georges M (2004) Genetic and functional confirmation of the causality of the DGAT1 K232A quantitative trait nucleotide in affecting milk yield and composition. Proc Natl Acad Sci USA 101:2398–2403.

4. Sutton JD (1989) Altering Milk Composition by Feeding. J Dairy Sci 72:2801–2814.

5. Benchaar C, Petit HV, Berthiaume R, Whyte TD, Chouinard PY (2006) Effects of Addition of Essential Oils and Monensin Premix on Digestion, Ruminal Fermentation, Milk Production, and Milk Composition in Dairy Cows. J Dairy Sci 89:4352–4364.

6. Perry FG, Macleod GK (1968) Effects of Feeding Raw Soybeans on Rumen Metabolism and Milk Composition of Dairy Cows. J Dairy Sci 51:1233–1238.

7. Mattos R, Staples CR, Arteche A, Wiltbank MC, Diaz FJ, Jenkins TC, Thatcher WW (2004) The Effects of Feeding Fish Oil on Uterine Secretion of PGF2α, Milk Composition, and Metabolic Status of Periparturient Holstein Cows. J Dairy Sci 87:921–932.

8. Grisart B, Coppieters W, Farnir F, Karim L, Ford C, Berzi P, Cambisano N, Mni M, Reid S, Simon P, Spelman R, Georges M, Snell R (2002) Positional Candidate Cloning of a QTL in Dairy Cattle: Identification of a Missense Mutation in the Bovine DGAT1 Gene with Major Effect on Milk Yield and Composition. Gen Res 12:222–231.

9. Cheng D, Iqbal J, Devenny J, Chu C-H, Chen L, Dong J, Seethala R, Keim WJ, Azzara AV, Lawrence RM, Pelleymounter MA, Hussain MM (2008) Acylation of Acylglycerols by Acyl Coenzyme A:Diacylglycerol Acyltransferase 1 (DGAT1) - Functional importance of DGAT1 in the intestinal fat absorption. J Biol Chem 283:29802–29811.

10. Mach N, Goselink RMA, van Baal J, Kruijt L, van Vuuren AM, Smits MA (2013) Relationship between milk fatty acid composition and the expression of lipogenic genes in the mammary gland of dairy cows. Livest Sci 151:92–96.

11. Clarke SD (2000) Polyunsaturated fatty acid regulation of gene transcription: a mechanism to improve energy balance and insulin resistance. Br J Nutrit 83(S1):S59–S66.

12. Lapillonne A, Clarke S, Heird WC (2004) Polyunsaturated fatty acids and gene expression. Curr Opin Clin Nutr Metab - Lipid metab Ther 7:151–156.

13. Mach N, Jacobs AAA, Kruijt L, van Baal J, Smits MA (2011). Alteration of gene expression in mammary gland tissue of dairy cows in response to dietary unsaturated fatty acids. Animal 5:1217–1230.

14. Kliewer SA, Sundseth SS, Jones SA, Brown PJ, Wisely GB, Koble CS, Devchand P, Wahli W, Willson TM, Lenhard JM, Lehmann JM 1997. Fatty acids and eicosanoids regulate gene expression through direct interactions with peroxisome proliferator-activated receptors α and γ. Proc Natl Acad Sci USA 94:4318–4223

15. West DB, Delany JP, Camet PM, Blohm F, Truett AA, Scimeca J (1998) Effects of conjugated linoleic acid on body fat and energy metabolism in the mouse. Am J Physiol – Regul Integr Comp Physiol 275:R667–R672.

16. Weljie AM, Newton J, Mercier P, Carlson E, Slupsky CM (2006) Targeted Profiling:Quantitative Analysis of 1H NMR Metabolomics Data. Anal Chem 78:4430–4442.

17. Wishart DS (2008) Quantitative metabolomics using NMR. Trend Anal Chem 27:228– 237.

18. Sundekilde UK, Frederiksen PD, Clausen MR, Larsen LB, Bertram HC (2011) Relationship between the Metabolite Profile and Technological Properties of Bovine Milk from Two Dairy Breeds Elucidated by NMR-Based Metabolomics. J Agric Food Chem 59:7360–7367.

19. Sundekilde UK, Larsen LB, Bertram HC (2013) NMR-Based Milk Metabolomics. Metabolites 3:204–222.

20. Mach N, van Baal J, Kruijt L, Jacobs A, Smits MA (2011). Dietary unsaturated fatty acids affect the mammary gland integrity and health in lactating dairy cows. BioMed Central Proc 5 (Suppl. 4), S35. http://www.biomedcentral.com/1753-6561/5/S4/S35.

21. Mach N, Blum Y, Bannink A, Causeur D, Houee-Bigot M, Lagarrigue S, Smits MA (2012) Pleiotropic effects of polymorphism of the gene diacylglycerol-Otransferase 1 (DGAT1) in the mammary gland tissue of dairy cows. J Dairy Sci 95:4989–5000.

22. Mach N, Zom RLG, Widjaja HCA, van Wikselaar PG, Weurding RE, Goselink RMA, van Baal J, Smits MA, van Vuuren AM (2013). Dietary effects of linseed on fatty acid composition of milk and on liver, adipose and mammary gland metabolism of periparturient dairy cows. J Anim Physiol Anim Nutr 97:89–104.

23. Jacobs AAA, van Baal J, Smits MA, Taweel HZH, Hendriks WH, van Vuuren AM, Dijkstra J (2011) Effects of feeding rapeseed oil, soybean oil, or linseed oil on stearoyl-CoAdesaturase expression in the mammary gland of dairy cows. J Dairy Sci 94:874–887.

24. Schennink A, Stoop WM, Visker MHPW, Heck JML., Bovenhuis H, van der Poel JJ, van Valenberg HJF, van Arendonk JAM (2007) DGAT1 underlies large genetic variation in milk-fat composition of dairy cows. Anim Genet 38:467–473.

25. Klein MS, Almstetter MF, Schlamberger G, Nürnberger N, Dettmer K, Oefner PJ, Meyer HHD, Wiedemann S, Gronwald W (2010) Nuclear magnetic resonance and mass spectrometry-based milk metabolomics in dairy cows during early and late lactation. J Dairy Sci 93:1539–1550.

26. Lu J, Boeren S, van Hooijdonk T, Vervoort J, Hettinga K (2015) Effect of the DGAT1 K232A genotype of dairy cows on the milk metabolome and proteome. J Dairy Sci 98: 3460–3469.

27. Kanehisa M, Goto S (2000) KEGG: Kyoto Encyclopedia of Genes and Genomes. Nucl Acid Res 28:27–30.

28. GeneOntology Consortium (2004) The Gene Ontology (GO) database and informatics resource. Nucl Acid Res 32(suppl 1):D258–D261.

29. Streeper RS, Koliwad SK, Villanueva CJ, Farese RV (2006) Effects of DGAT1 deficiency on energy and glucose metabolism are independent of adiponectin. Am J Physiol Endocrinol Metab 291:E388–E394.

30. Sorensen B, Kühn C, Teuscher F, Schneider F, Weselake R, Wegner J (2006) Diacylglycerol acyltransferase (DGAT) activity in relation to muscle fat content and DGAT1 genotype in two different breeds of Bos Taurus. Arch Anim Breed 49:351–356.

31. Scriver CR, Gregory DM, Sovetts D, Tissenbaum G (1985) Normal plasma free amino acid values in adults: The influence of some common physiological variables. Metab 34:868–873.

32. Taschuk R, Griebel PJ (2012) Commensal microbiome effects on mucosal immune system development in the ruminant gastrointestinal tract. Anim Health Res Rev 13:129– 141.

33. De Menezes AB, Lewis E, O’Donovan M, O’Neill BF, Clipson N, Doyle EM (2011) Microbiome analysis of dairy cows fed pasture or total mixed ration diets. Fed Eur Microbiol Socs Microbiol Ecol 78:256–265.

34. Fernando SC, Purvis II HT, Najar FZ, Sukharnikov LO, Krehbiel CR, Nagaraja TG, Roe BA, DeSilva U (2010) Rumen Microbial Population Dynamics during Adaptation to a High-Grain Diet. Appl Environm Microbiol 76:7482–7490.

